# Transcriptomic Meta-analysis Identifies Long Non-Coding RNAs Mediating Zika’s Oncolytic Impact in Glioblastoma Multiforme

**DOI:** 10.1101/2024.08.04.605859

**Authors:** Youssef A. Kousa, Shriya Singh, Allison Horvath, Federica Tomasso, Javad Nazarian, Lisa Henderson, Tamer A. Mansour

## Abstract

Glioblastoma multiforme (GBM) is an aggressive and lethal form of brain cancer with few effective treatments. In this context, Zika virus has emerged as a promising therapeutic agent due to its ability to selectively infect and kill GBM cells. To elucidate these mechanisms and expand the landscape of oncolytic virotherapy, we pursued a transcriptomic meta-analysis comparing the molecular signatures of Zika infection in GBM and neuroblastoma (NBM). Over-representation analysis of dysregulated coding genes showed significant enrichment of tumor necrosis factor (TNF), NF-κB, and p53 signaling pathways. A refined list of long non-coding RNAs consistently dysregulated in Zika-infected GBMs was also developed. Functional review of these candidates revealed their potential regulatory role in Zika-mediated oncolysis. We performed validation of the less-researched targets in adult and pediatric GBM cell lines and found significant differential regulation, as predicted. Altogether, our results provide novel insights into the molecular mechanisms underlying the effect of Zika on GBM. We highlight potential therapeutic targets that could be further interrogated to improve the efficacy of tumor cell death and the utility of Zika as an adjuvant virotherapy for GBM and other related cancers.

## INTRODUCTION

Primary malignant brain tumors affect over 300,000 individuals each year, and most are incurable.^1^ High grade gliomas (HGG) and glioblastoma multiforme (GBM) are aggressive cancers, causing significantly high morbidity and mortality in affected children and adults. A combination of chemo- and radiotherapy is the current standard of care, although success is limited due to high rates of resistance and recurrence.^2,3^ As a result, there is a gap in effective treatment and management of GBM, resulting in a median survival of 15 months following diagnosis.^4-6^ This is especially critical given concern for the global rise of GBM’s incidence rate.^5,6^ Considering such limitations, there is urgent need for novel adjuvant therapies to improve treatment and prognosis for patients with GBM.

Oncolytic viruses are emerging as a potential adjuvant therapy for GBM. In theory, such viruses selectively induce cell death by targeting proliferating tumor cells, while sparing quiescent healthy cells.^7^ Among neurotropic viruses, Zika virus has emerged as a potential neuro-oncolytic agent. Zika garnered substantial attention after recent outbreaks in South America, the Caribbean, Central America, Africa, and Asia. In 2016, prenatal infection was associated with developmental brain injury and microcephaly, causing the World Health Organization (WHO) to declare the pandemic a global health emergency of international concern.^8,9^ Zika induces prenatal brain injury due to a unique combination of neurotropism and proclivity for proliferating cells.^10,11^

While injurious to the prenatal brain, these features have added interest in Zika as an oncolytic virotherapy.^12,13^ Phenotypic and transcriptional similarities between neural and glioblastoma stem cells have focused special attention on the use of Zika virus in GBM.^14^ In fact, several proteins needed for Zika to bind (SOX2-Integrins) and replicate (MSI1) in cells are expressed at higher levels in glioblastoma stem cells.^15,16^ Consistently, Zhu *et al*. 2017 revealed that inoculation of a mouse-adapted Zika strain in xenografted GBM cells led to inhibited growth of the tumor cells and significantly improved survival in mice.^13^ In addition to GBM, certain neuroblastoma (NBM) cell lines are highly permissive to Zika infection, resulting in greater cellular cytotoxicity and reduced tumor cell viability.^12,17^

In contrast to GBM, which originate from glial cells, NBM is predominantly a pediatric cancer that arises from immature neuroblasts in the developing sympathetic nervous system.^18,19^ NBM is also markedly distinct in its molecular profile, clinical presentation, and outcomes. Despite the differences, NBM and GBM are vulnerable to Zika infection and oncolysis. It is not clear if shared or distinct mechanisms mediate the impact on different tumor types.

Three studies have characterized the effect of Zika infection on GBM and NBM. Zhu *et al*. and Bonenfant *et al*. 2020 performed transcriptomic studies to explain the molecular changes underlying GBM and NBM respectively but with limited statistical power and exclusive focus on coding genes.^13,17^ More recently, Bulstrode *et al*. 2022 used transcriptomic analysis for a large cohort of GBM tumor specimens to assess the effect of interferon secretion on malignant neural progenitor cells without much focus on the oncolytic effect of the virus in GBM.^12^ A meta-analysis of these transcriptomic profiles has not been performed and is needed for many reasons, including: 1) Eliminating the batch effects of individual studies; 2) Evaluating for differential expression of non-coding genes; and 3) Minimizing the high false positive rates associated with low-powered omic studies.^20^

Toward deepening our understanding of Zika’s potential for bridging treatment gaps, we systematically evaluated the transcriptional profiles of Zika-infected GBM. Harmonizing four transcriptional datasets from three publications, we aimed to uncover molecular mechanisms mediating Zika’s neuro-oncolytic impact. Using NBM as an outgroup, we identified genes and pathways targeted by Zika virus among GBM studies. We observed that Zika dysregulates several canonical cancer pathways in mediating cell death, including tumor necrosis factor (TNF), NF-κB, and p53 signaling. Scrutiny of dysregulated non-coding genes revealed unexpected novel targets, including long non-coding RNAs (lncRNAs), which were transcriptionally validated. Together, these data suggest mechanistic insights and avenues for developing neuro-oncolytic virotherapies.

## METHODS

### Data

Illumina sequencing reads for Zika-infected GBM samples and their matched controls were downloaded from SRA for PRJNA399336 (50SE) and PRJNA739733 (50PE) (Table S1). Similarly, infected NBM samples and controls were downloaded for PRJNA630088 (124PE).

### Inclusion/exclusion criteria

Studies linked to publicly available data in Gene Expression Omnibus (GEO) Datasets were searched to identify transcriptomic datasets that investigated neural tumor response to Zika infection. Such studies are limited in the literature and were evaluated against a standardized set of inclusion criteria, which included: RNA-sequencing of total transcriptomic changes and a study design composed of at least two biological replicates in the Zika-infected and mock-infected groups.

### Reference transcriptome

RefSeq gene annotation showed better quantification accuracy compared to Ensembl annotation. However, recent expansion in the number of gene models, specifically those with a smaller size, had a detrimental effect.^21^ Therefore, the RefSeq release v.110 of gene annotations for Homo sapiens were downloaded from NCBI and filtered to exclude transcripts smaller than 250bp. To adjust for the possible compositional change of gene expression because of viral gene expression in infected samples, seven viral genomes were added to the reference as multiple isoforms of one gene. All complete viral genomes were downloaded from the Nucleotide database of NCBI (n=328). The genomes were clustered into groups with a minimum of 95% sequence similarity by cd-hit-est version 4.6.^22^ The longest sequence in each group was selected as a representative.

### Differential expression analysis

Raw FASTQ sequences were trimmed by Trimmomatic v.39 to remove adaptors and low-quality sequences.^23^ High quality reads were mapped to the reference transcriptome by bowtie2 v.2.3.4.3.^24^ BAM files were used for quantification of transcript abundance by salmon v.1.9.0 with correction for fragment GC content bias.^25^ R v.4.2.1 (2022-06-23) was used for subsequent differential expression analysis.^26^ Transcript-level estimates were corrected for the transcript length and summarized for gene level analysis using tximeta package v.1.14.1.^27^ The DESeq2 package v.1.36.0 was used for differential expression testing using a negative binomial GLM fitting and Wald statistics.^28^ The model was adjusted for the biosource in PRJNA739733. Differentially expressed genes were defined by false discovery rate (FDR) less than 0.05. Among the datasets, differentially expressed genes were identified and divided into coding and non-coding groups by home-made scripts.

### Gene expression clustering analysis

After summarization of the counts per gene, the data was normalized for the library size and its variance was stabilized using the variance stabilizing transformation function in the DESeq2 R package. The counts from the Bulstrode *et al*. experiment were adjusted for the sample-source batch effect using the limma R package. The counts of the three studied experiments were then combined and adjusted for the study batch effect using the same function. The adjusted count matrix of the differentially expressed genes was used to cluster the differentially expressed genes and assess similarity between samples used in a heatmap and a PCA plot by the pheatmap and plotPCA functions from the pheatmap and DESeq2 R package.

### Functional enrichment analysis

WebGestalt^29^ was used to test protein-coding differentially expressed genes for pathway enrichment using over-representation testing in KEGG pathways, DrugBank, miRNA targets, and transcription factor targets from the MSigDB database using known protein-coding genes as a background. For each analysis, the minimum and maximum number of genes for a category were set to five and 2000, respectively. The categories were first ranked based on FDR and then the top 10 most significant categories were selected. This was done separately for up- and down-regulated categories. The software used Benjamini-Hochberg adjustment to adjust p values as it tested multiple gene sets simultaneously. In the analysis of differentially regulated lncRNAs, we reviewed NCBI’s GeneRIF database, and findings pertaining to GBM, NBM, gliomas, and other tumors, were prioritized, as available. Findings summarized in Fig. 6 was created with BioRender.com.

### Glioblastoma cell cultures

U87 cells (ATCC, USA) were cultured in Eagle’s Minimum Essential Medium (EMEM, ATCC®) supplemented with 10% Fetal Bovine Serum (FBS, GibcoTM) and 1% Penicillin/Streptomycin (Sigma-Aldrich). GBM110 cell line (BTRL, USA) was cultured in NeuroCult NS-A Basal Medium-Human (Stem Cell Technologies) supplemented with NeuroCult NS-A Proliferation Supplements-Human (Stem Cell Technologies), 10 ng/ml bFGF (PeproTech), 10 ng/ml EGF (PeproTech), and 1% Penicillin/Streptomycin. For GBM110, plates were coated with 1 µg/µl Laminin from Engelbreth-Holm-Swarm murine sarcoma basement membrane (Sigma-Aldrich) to allow cell adherence. Cells were maintained in a humidified incubator at 37°C and 5% CO_2_.

### Zika titration

Zika French Polynesian strain (Nath Lab, NINDS) was expanded in Vero cells (ATCC) by inoculating cells at an MOI of 0.01 in FBS free-DMEM, high glucose, GlutaMAX™ Medium (Gibco™) and incubating for 72 h at 37°C and 5% CO_2_. The supernatant containing the virus was mixed with 1X Sucrose-Phosphate-Glutamate, then filtered and stored at −80°C. Viral titers were quantified by plaque forming assay. Viral stocks were serially diluted (10^-1^ to 10^-6^) and 100 µl of viral dilution were added to Vero cells in monolayer in a 24 well plate. After 1 h, 900 ul of overlay media (1% Methylcellulose, 20% PBS, 2% FBS, DMEM with GlutaMAX™, pH 7.5) was added to each well and cells were incubated for 5 days at 37°C. Cells were fixed in 4% Paraformaldehyde (PFA) and stained with crystal violet dye. Foci were quantified to determine viral titer.

### Zika infection

We virally infected U87 (adult) and GBM110 (pediatric) cells in 6 well-plates (1×10^5^ cells/well). Cells were infected at 80% confluency using an MOI of three (GBM110; Mock N = 5, Zika-infected N = 6) or at an MOI of 0.5 (U87; Mock N = 6, Zika-infected N = 5), in FBS and Pen/Strep free-medium. After four hours, virus containing-medium was removed and replaced with standard culture media. Cells were incubated for 48h at 37°C and 5% CO_2_.

### RT-qPCR

Cell pellets were collected at 48 hours post-infection. RNA was extracted using RNeasy Plus Mini Kit (Qiagen), following manufacturer instructions. All RNA was normalized to 25ng/uL and cDNA was produced with SuperScript™ III First-Strand Synthesis SuperMix for qRT-PCR from Invitrogen. cDNA was quantified using Fast SYBR™ Green Master Mix from Applied Biosystems on QuantStudio 7 Flex Real-Time PCR System with the primers indicated in Table S2 (LINC03032, TIPARP-AS1, SH3RF3-AS1, EMC3-AS1, HCG18, MELTF-AS1). 5 mock and 6 infected biological replicates were used to analyze U87 cells. 6 mock and 5 infected replicates were used to analyze GBM 110 cells. Genes were normalized to beta-actin. Data analysis and figures were made using Prism. An unpaired, two-tailed student’s t-test was used in determining significance between mock and Zika-infected samples.

## RESULTS

We compared differentially expressed genes among transcriptional profiles obtained from Zika-infected GBM and NBM studies. As expected, the analysis revealed significant differences among gene expression profiles between the GBM and NBM studies; there was more overlap within the two GBM studies and the two NBM studies (Fig. 1). There were very few genes differentially regulated among all four datasets, especially for long non-coding RNA (Fig. 1C, D).

**Figure 1.**
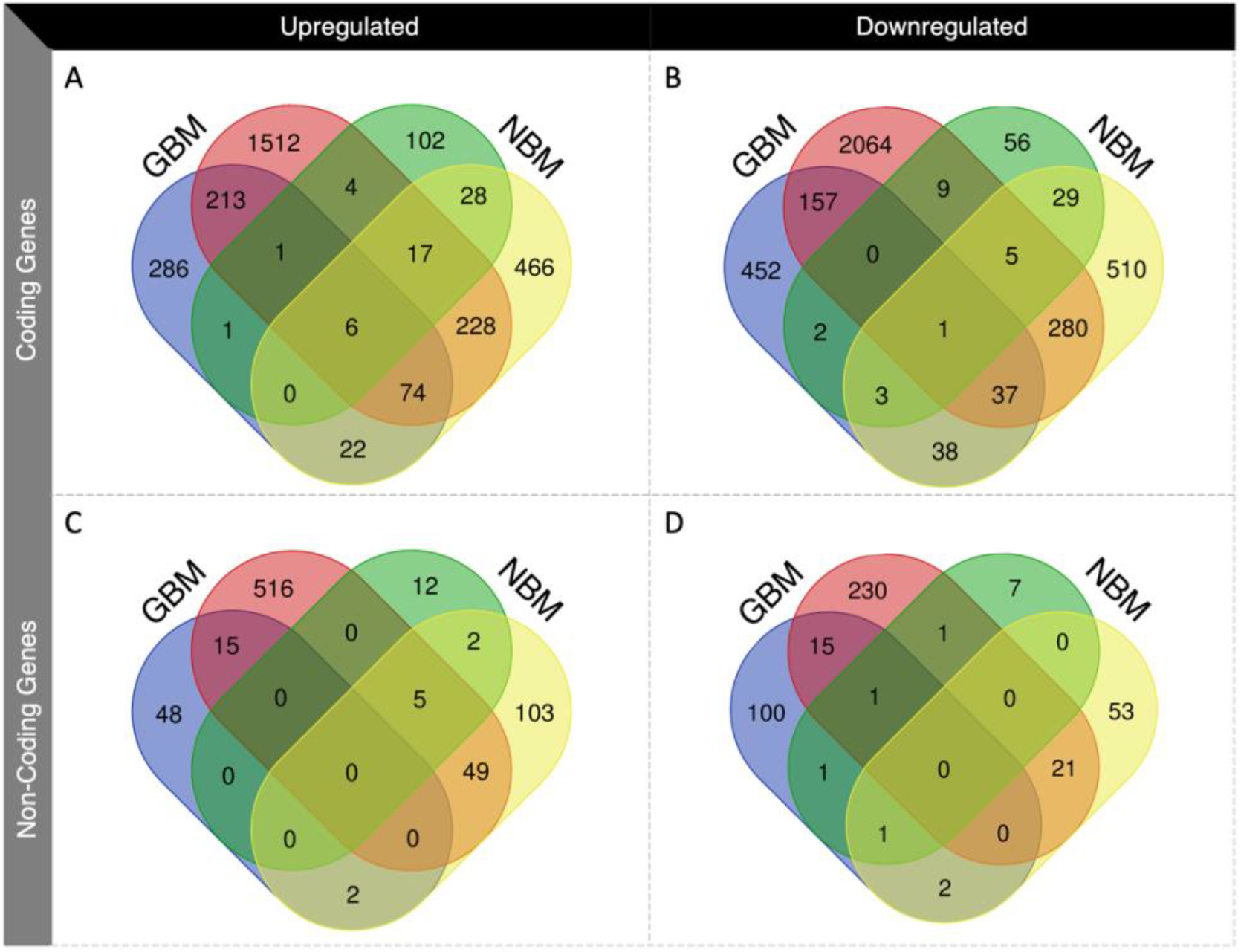
Comparative analysis of GBM and NBM studies. GBM studies correspond with blue (Bulstrode *et al*. 2022) and red (Zhu *et al*. 2017) highlighting while NBM studies correspond with green and yellow (Bonenfant *et al*. 2020) highlighting. (A and B) Differentially expressed protein-coding genes across the studies that were (A) upregulated or (B) downregulated after Zika-infection. (C and D) Differentially expressed non-coding genes that were (C) upregulated or (D) downregulated after Zika-infection.

Principal component analysis (PCA) evaluated the degree of variability between control and Zika-infected GBM and NBM between different viral strains (Fig. 2). As expected, the differentially expressed coding and non-coding genes in GBM indicate clear separation of the Zika-infected samples from their matching controls. Among the coding genes, uninfected and infected NBM profiles are more closely related to uninfected GBM (Fig. 2A). However, among non-coding genes, there is a clear segregation between GBM and NBM, and between uninfected and infected samples (Fig. 2B). This is reflective of the unique nature of the differentially expressed genes in GBM after Zika-infection, irrespective of viral strain. Heatmaps were also generated for both coding and non-coding differentially expressed genes (Fig. S1).

**Figure 2.**
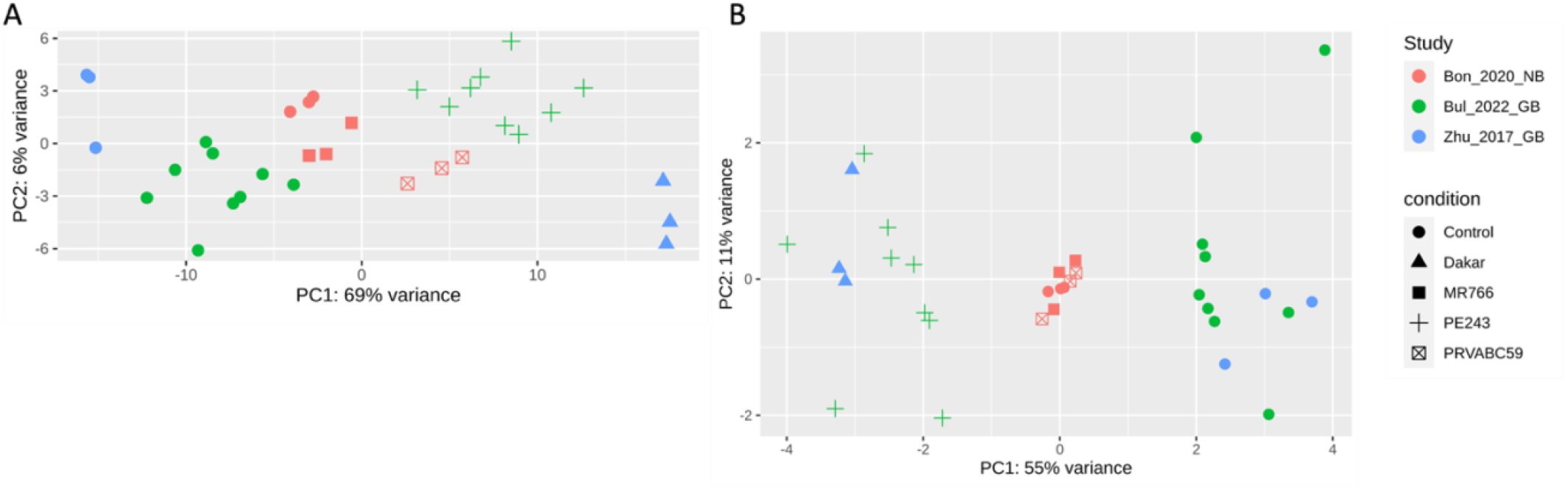
PCA of batch-corrected gene expression. Principal component analysis (PCA) assessed variance within differentially expressed coding (A) and non-coding (B) genes between control and Zika-infected (Dakar/MR766/PE243/PRVABC59-infected) cells.

Meta-analysis of the two GBM studies identified a consensus of 294 upregulated and 195 downregulated protein-coding genes (Fig. 1A, B). Both lists of genes were tested for functional enrichment by performing an over-representation analysis (ORA) for molecular pathways, pharmaceutical agents, miRNA targets, and transcription factor binding sites. KEGG pathway analysis identified upregulation of genes driving TNF, NF-κB, and p53 signaling (Fig. 3A). ORA against pharmaceutical agents in the DrugBank database indicated significant enrichment of upregulated genes for the molecular targets of andrographolide, a natural compound with anti-oncogenic properties (Fig. 3B).^30-32^ While enrichment analysis against miRNA targets did not reach FDR threshold, the top two hits, MIR-502 and MIR-362, both have known tumor suppressor function (Fig. 2C).^33-35^ Finally, upregulated genes were highly enriched in several transcription factor binding sites for NF-κB signaling, which plays a key role in mediating the relationship between cancer and inflammation, and CREB, which is an established proto-oncogene (Fig. 3D).^36-38^ Similar analyses of downregulated protein-coding genes in GBM were not statistically significant (Fig. 3E, F, G, H).

**Figure 3.**
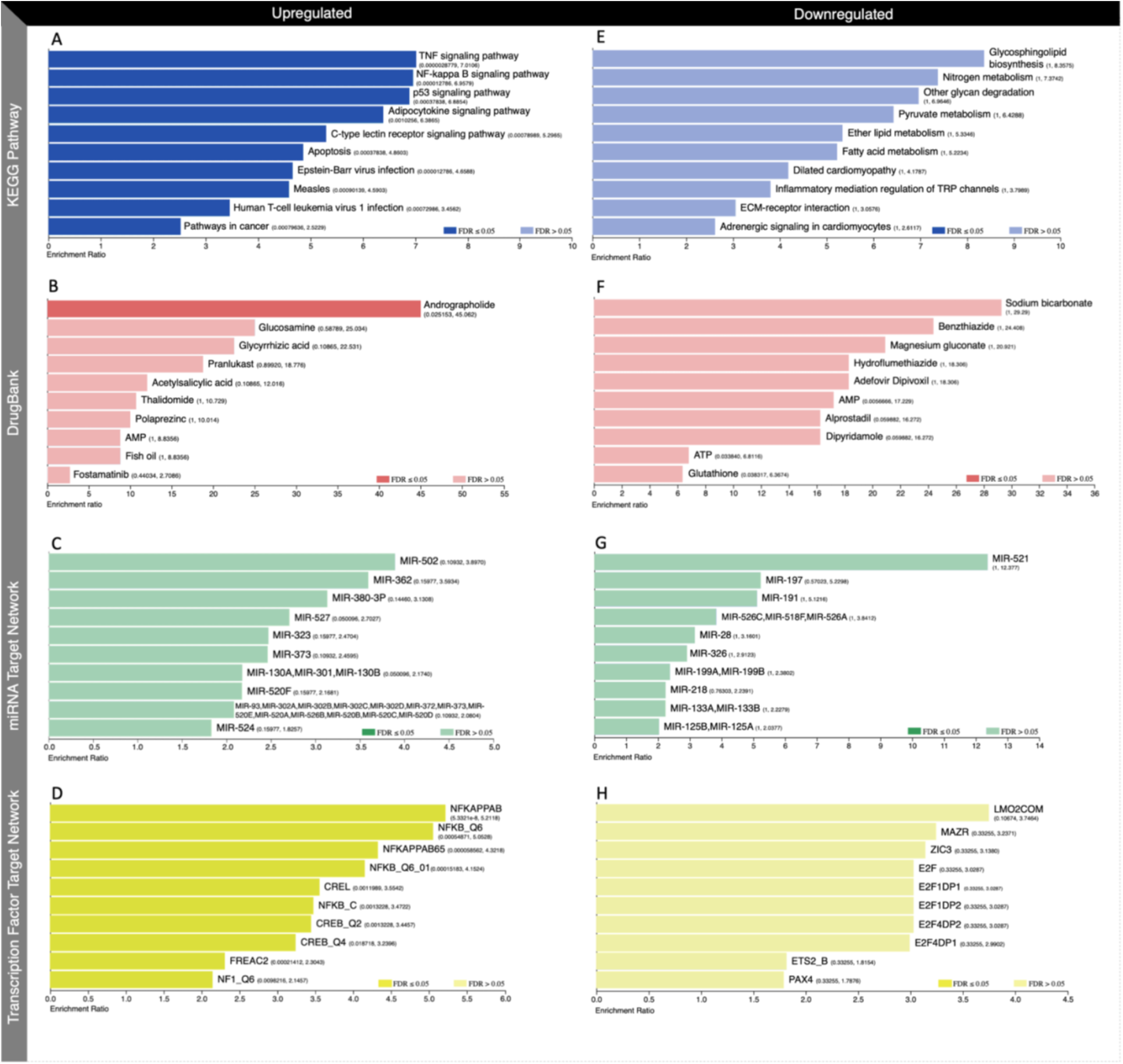
Over-representation analysis of protein-coding genes dysregulated in GBM. The bar graphs show the functional enrichment analysis of upregulated (A, B, C, D) and downregulated (E, F, G, H) genes in KEGG pathways (A, E), DrugBank (B, F), miRNA targets (C, G), and (D, H) transcription factor targets. Significance was defined by false discovery rate (FDR) ≤ 0.05. The bars are annotated by p-values followed by enrichment ratios.

Next, we conducted a meta-analysis to combine the GBM and NBM studies and evaluate if shared or distinct mechanisms mediate Zika’s impact on different tumors. We included coding genes differentially expressed in at least three out of the four datasets (Fig. S2A). As might be expected, this analysis resulted in shorter lists of differentially expressed genes that consisted of 98 upregulated and 46 downregulated protein-coding genes (Fig. 1A, B). As in the pathway analysis between the GBM tumors, upregulated genes were enriched in pathways related to TNF signaling, NF-κB signaling, and apoptosis (Fig. 4A). While andrographolide remained the most enriched drug for the upregulated genes, it failed to meet the FDR threshold (Fig. 4B). Instead, acetylsalicylic acid also emerged as a drug candidate. Acetylsalicylic acid is known to inhibit GBM angiogenesis, proliferation, and motility, and has been proposed to produce a similar effect in NBM.^39-42^ None of the miRNA targets reached threshold, but MIR-202, one of the top three enriched targets, has been shown to act as a tumor suppressor in both GBM and NBM (Fig. 4C).^43-45^ Upregulated genes continued to be highly enriched in NF-κB and CREB transcription factor targets, while STAT3’s enrichment was nearly four-fold greater than the second predicted target (Fig. 4D).

**Figure 4.**
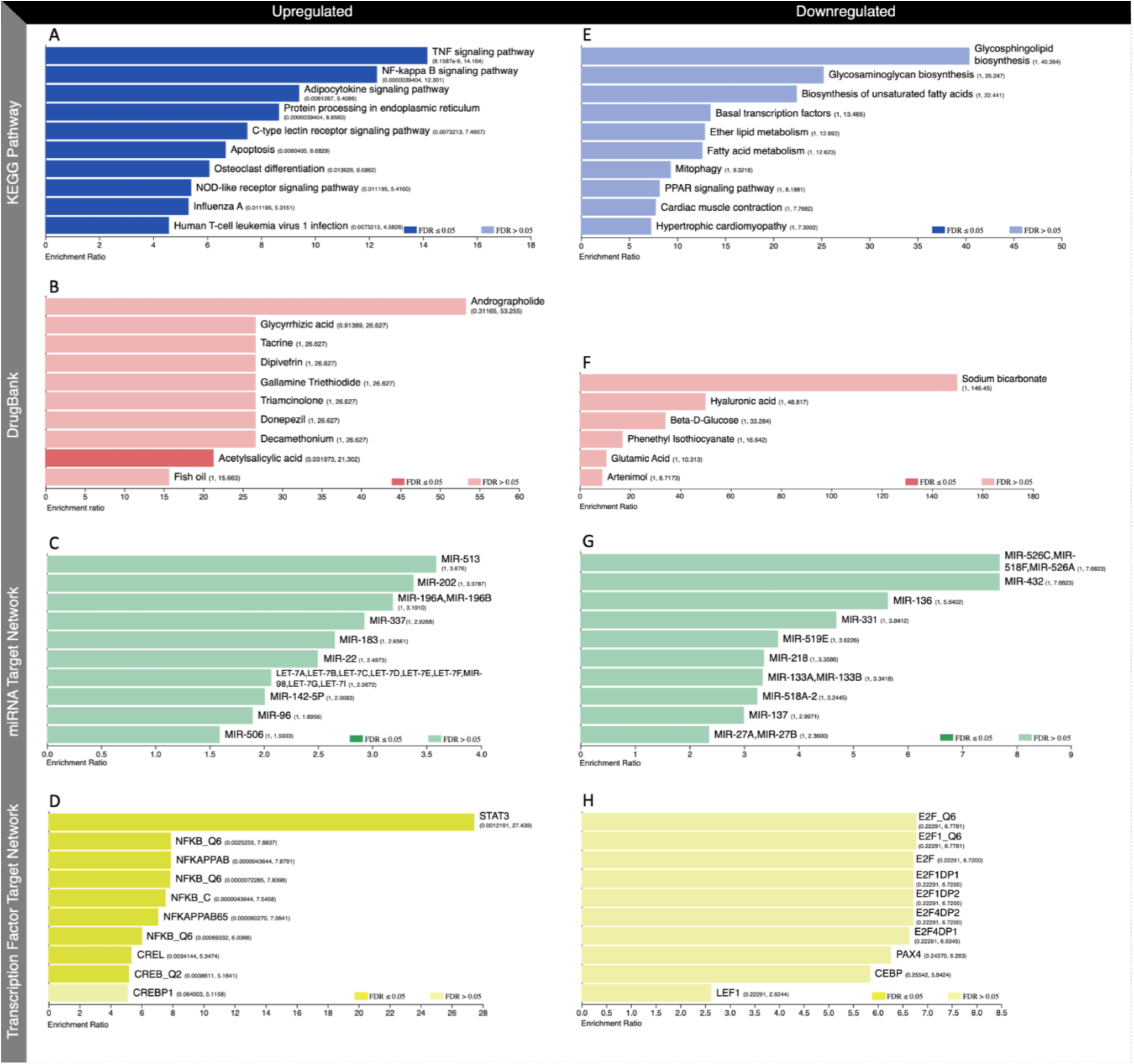
Over-representation analysis of protein-coding genes dysregulated in the combined GBM and NBM analysis. The bar graphs show the functional enrichment analysis of upregulated (A, B, C, D) and downregulated (E, F, G, H) genes in KEGG pathways (A, E), DrugBank (B, F), miRNA targets (C, G), and (D, H) transcription factor targets. Significance was defined by false discovery rate (FDR) ≤ 0.05. The bars are annotated by p-values followed by enrichment ratios.

Enrichment of downregulated genes mirrored the results from the analysis between the two GBM data sets for the KEGG pathways and DrugBank analysis. Although not meeting FDR threshold, downregulated genes in the GBM and combined GBM/NBM analysis were enriched for lipid and fatty acid metabolic pathways and sodium bicarbonate treatment (Fig. 3E, F and 4E, F). Interestingly, we noted overlap among miRNA targets and the E2F transcription factor binding sites (Fig. 4G, H).

The function and roles of non-coding RNA continues to emerge with next generation sequencing. Our analysis identified 31 consistently dysregulated long non-coding RNAs (15 upregulated and 16 downregulated) between the two GBM studies (Fig. 1C, D; Fig. S2B). After excluding uncharacterized lncRNAs and pseudogenes, we identified 12 genes of interest. Each of these were evaluated for possible or known functions, in comparison with previously reported gene expression changes (Table 1). Transcriptomic data showed that five of seven lncRNAs previously identified in oncogenesis had expression changes predicted in mediating tumor cell death (for instance, downregulation of an oncogene).

**Table 1.** Summary of likely functional lncRNA candidates. Expression indicates the type of dysregulation detected in our meta-analysis.

| Gene | Tumor Type | Literature Review<br>Possible / Known Function | Predicted<br>Expression in GBM | Ref |
| --- | --- | --- | --- | --- |
| NR2F1-AS1 | Neuroblastoma | Tumor Suppressor: targets miR-493/TRIM2 axis<br>Upregulation modulated to suppress cell proliferation and migration, while increasing apoptosis | Upregulated | 52 |
| LINC03032 | - | - | Upregulated | - |
| TIPARP-AS1 | Breast Cancer | Regulates TIPAP-mediated AHR Signaling | Upregulated | 53 |
| SH3RF3-AS1 | - | - | Upregulated | - |
| LINC01783 | Tongue Squamous Cell Carcinoma | Oncogene: targets miR-199b-5p<br>Upregulation promoted cell proliferation and metastasis | Upregulated | 54 |
| SNX10-AS1 | - | - | Downregulated | - |
| SLC9A3-AS1 | Nasopharyngeal Carcinoma | Oncogene: targets miR-486-5p/E2F6 axis<br>Loss of SLC9A3-AS1 reduced cell proliferation and metastasis, while inducing apoptosis in vitro; reduced tumor growth in vivo | Downregulated | 55 |
| EMC3-AS1 | - | - | Downregulated | - |
| HCG18 | Anaplastic Glioma | Tumor Suppressor/Protective Factor<br>Downregulation associated with increased tumor grade | Downregulated | 55 |
| MELTF-AS1 | Glioblastoma Multiforme | Oncogene: targets miR-485-5p/MMP14 axis<br>Downregulation suppresses tumor growth and metastasis, while inducing apoptosis; silencing represses tumor growth in vivo | Downregulated | 57 |
| PANTR1 | Glioblastoma Multiforme | Oncogene: targets POU3F3<br>Loss of PANTR1 decreased proliferation, colony formation, and viability | Downregulated | 58 |
|  |  | Silencing decreased expression of pro-angiogenesis factors (bFGF, VEGFA, bFGFR, and Angio) |  | 59 |
| MSH5-SAPCD1 | - | - | Downregulated | - |

We selected three previously associated and three novel lncRNA for further validation (six of 12 identified). Validation was performed with direct measurement of transcriptional levels in two Zika-infected GBM cell lines: one adult (U87) (Fig. 5A,B) and one pediatric (GBM110) (Fig. 5B,D). The selected MOIs allowed cell survival, while visibly altering cell morphology toward cell death (Fig. 5A,B).

**Figure 5.**
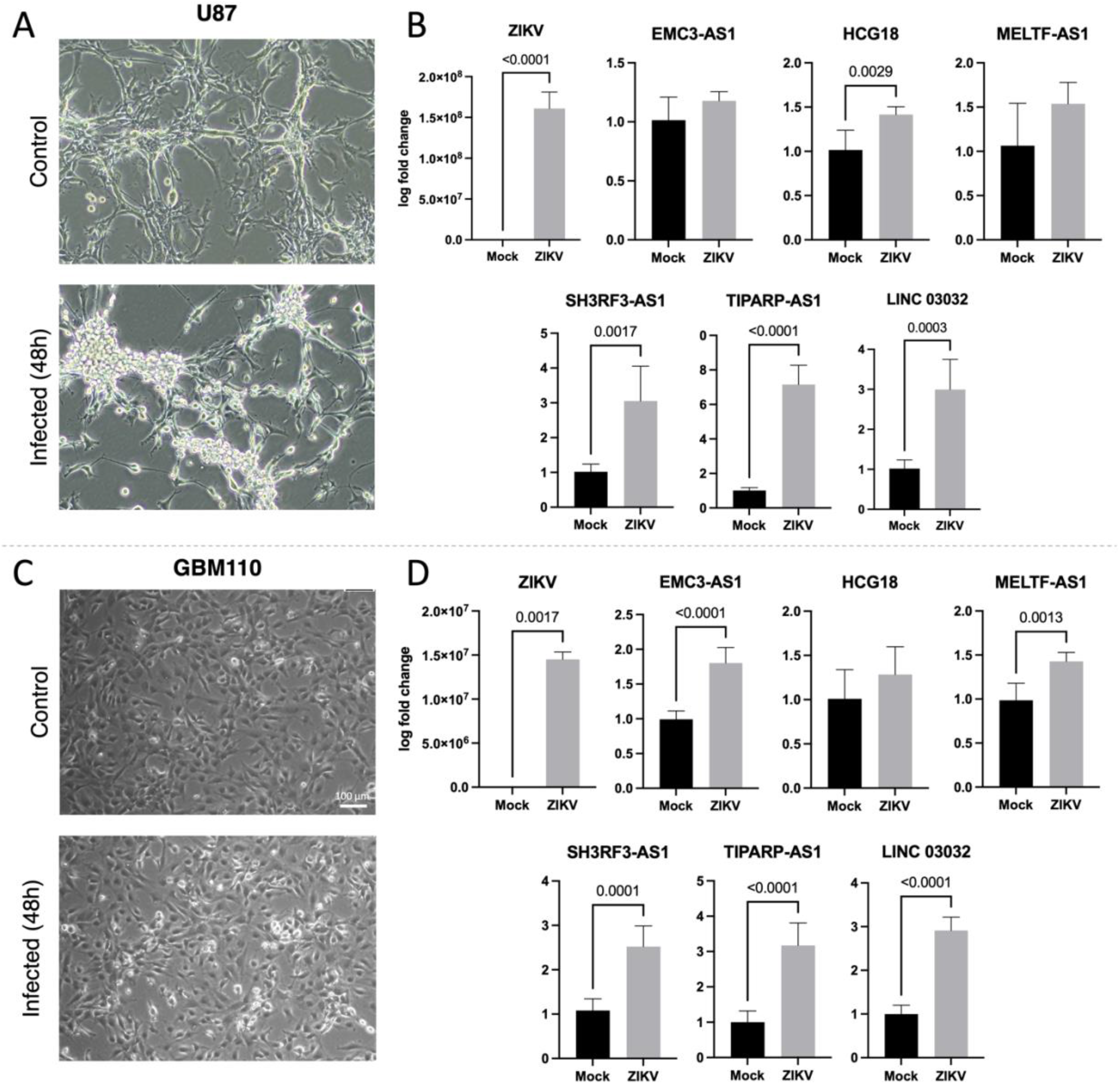
Transcriptionally validated lncRNA targets. Adult GBM cell line U87 (A) and pediatric cell line GBM110 (C) were infected at an MOI of 0.5 and 3, respectively. The cells were imaged 48 hpi after Zika-infection at 10x magnification. Scale bar is 100 microns (C). RT-qPR was completed using 5 mock/6 infected U87 biologicl replicates (B) and 6 mock/ 5 infected GBM 110 biological replicates (D). Transcriptional levels of six lncRNAs are measured by log fold change and p-values are indicated (B,D).

**Figure 6.**
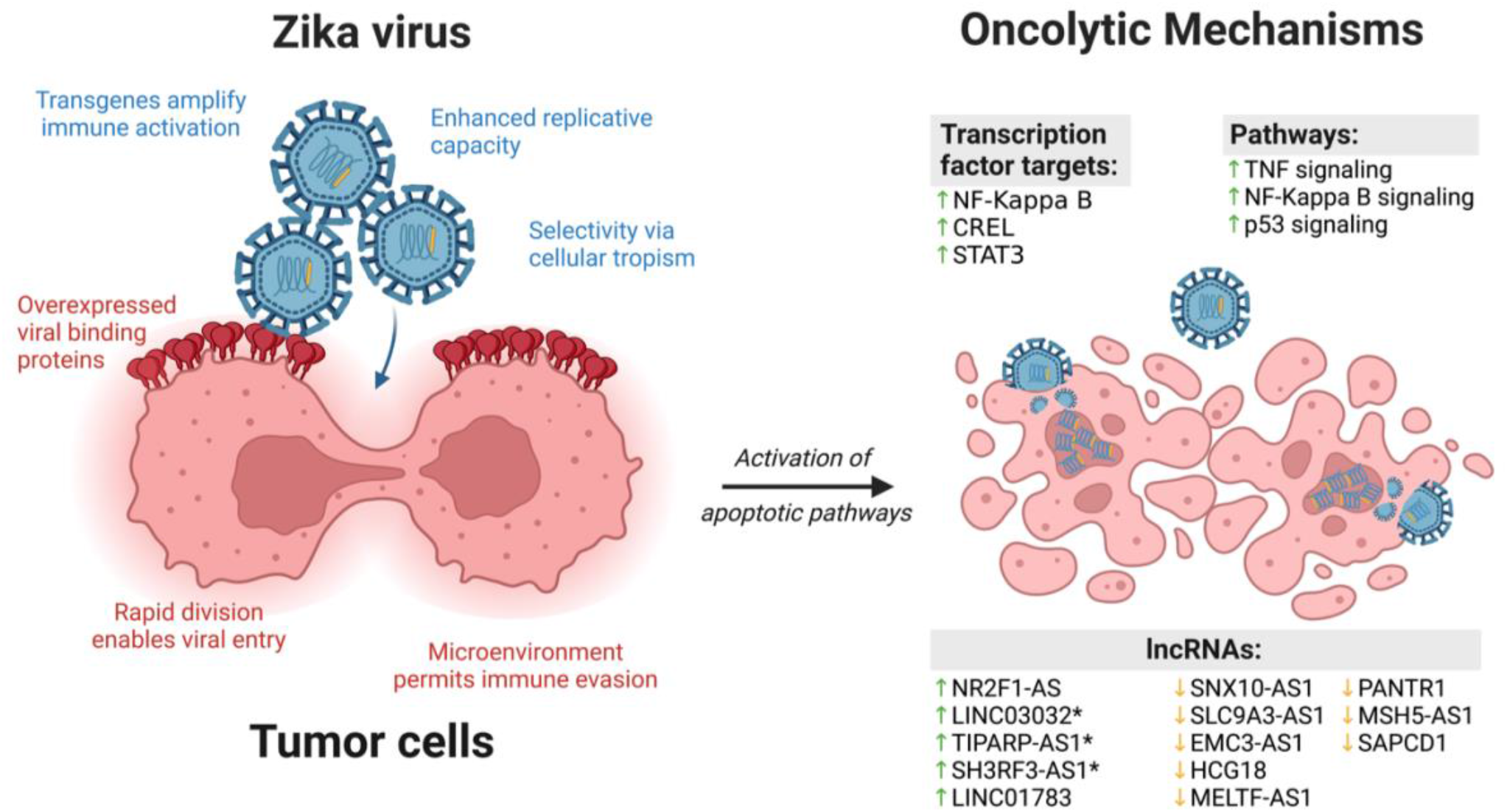
A working model summarizing how a combination of tumor susceptibilities and viral factors coalesce in mediating oncolytic impact. Viral-specific factors (blue), include: 1) modifying the viral genome by adding a transgene to regulate immune activation; 2) fine-tuning infectious dose or replication within the tumor cell; and 3) cellular targeting via innate tropism. Tumor-specific factors (red), include: 1) overexpressing viral binding proteins; 2) enabling rapid viral entry during cell division; and 3) evading immune surveillance as a byproduct of the tumor microenvironment. Our meta-analysis suggests a virally induced oncolytic impact mediated through differential regulation of select 1) transcription factors; 2) gene signaling pathways; and 3) lncRNAs. Asterisks for the three indicated genes indicate transcriptional validation in two cell lines

The three previously associated were MELTF-AS1 (GBM), TIPARP-AS1 (breast cancer), and HCG18 (anaplastic glioma), while the three novel lncRNA included SH3RF3-AS1, LINC03032, and EMC3-AS1. Notably, TIPARP-AS1, SH3RF3-AS1, and LINC03032 were all highly upregulated, as predicted, following Zika infection in both cell lines examined with significance p ≤ 0.01. In contrast, expression of EMC3-AS1, HCG18 and MELTF-AS1 were significantly altered in only one of the two lines (Fig. 5B,D).

## DISCUSSION

There is emerging interest in the use of Zika as a neuro-oncolytic agent in combination with existing treatment regimens for GBM and NBM. Several features of brain tumors make them especially vulnerable to virally induced cell death, including: 1) overexpression of membrane proteins promoting viral binding; 2) rapid cell division further enabling entry into the cell; 3) high metabolic and replicative activity dysregulating molecular pathways that drive viral replication; and 4) evasion of the host innate and adaptive immune response by tumor cells permitting the infection to persist.^46^ Features of neurotropic viruses can also be modified and exploited to enhance their therapeutic utility. Studies thus far have shown consistent, reproducible impact in Zika’s ability to result in tumor cell death through: 1) cellular tropism and selectivity; 2) replication capacity; 3) activation of apoptotic pathways; and 4) potentially modifying the viral genome with transgenes to regulate the immune response.^47^ As a result of these many features, there is a natural fit in the use of neurotropic viruses, especially Zika, in the treatment of GBM (Fig. 6).

We evaluated for shared genes and molecular networks that might be mediating the mechanisms in Zika’s oncolytic activity. We identified upregulation of genes driving canonical tumor pathways, including TNF, NF-κB, and p53 signaling pathways and a refined list of lncRNAs that could be involved in oncolytic impact (Fig. 6). Our results highlight additional roles of well-known pathways and identify novel molecular signatures mediating Zika-induced tumor cell death.

A noteworthy association to be explored is andrographolide, which induces cell cycle arrest and apoptosis and suppresses migration in GBM.^30-32^ Apoptosis is induced through p53 expression, which was also identified in our meta-analysis after Zika infection.^30^ Interestingly, andrographolide and Zika infection share downstream regulation of CREB, TNF, NFKB2, and NFKB1, although the direction of gene expression changes were dissimilar.^32,48^ The utility of andrographolide has not been explored in the context of Zika-mediated oncolysis. Our findings warrant further investigation in follow-up studies. Acetylsalicylic acid only rose to statistical significance as a candidate in the combined analysis, likely resulting from incorporation of the NBM datasets.

Involvement of inflammatory and immune pathways in viral-oncolysis is well-documented and was also highlighted by Zhu *et al*. and Bulstrode *et al*.^12,13^ Both studies found significant enrichment of interferon-stimulated genes, which are known for their role in antiviral responses and regulation of the immune system. The two studies also identified that Zika’s antitumor efficacy can be enhanced through inhibition of type 1 interferons, TYK2, or JAK/STAT signaling. The role of genes and pathways identified in this study has not been evaluated previously and warrant further research.

The comparative analysis between GBM and NBM (Fig. 1) highlighted important differences and suggest specificity in Zika’s neuro-oncolytic effect. For example, only 83 out of 1184 (7%) lncRNAs were shared among any of the four studies. This is consistent with known differences in cellular origins and molecular profiles between GBM and NBM. It is worth noting that one of the validated lncRNA genes associated to oncogenesis, TIPARP-AS1, is located immediately upstream of the *TIPARP* exon 1 in the opposite orientation, and its expression has been shown to inhibit expression of TIPARP/PARP-7^49^. TIPARP/PARP-7 is frequently overexpressed in many cancers and aids in immune evasion by negatively regulating the type I IFN response. Small molecule inhibitors of TIPARP/PARP7 induce regression of lung cancer xenografts and boost tumor-specific adaptive immune responses in mouse cancer cell models, highlighting the importance of this pathway in oncogenesis^50^. Similarly, Zika-induced upregulation of TIPARP-AS1 may lead to increased expression of type I IFNs which can directly induce cell cycle arrest and cytotoxicity within tumors.

Pooling the datasets for a meta-analysis also offered a unique opportunity to identify non-coding RNAs that could be driving the neuro-oncolytic effects. For example, the function of lncRNA SH3RF3-AS1 is not currently known but it was highly enriched in both infected GBM lines (p=0.0008 and p<0.0001 for U87 and GBM110, respectively). Interestingly, one study investigating the role of lncRNAs commonly dysregulated in hepatocellular carcinoma (HCC) revealed that expression of SH3RF3-AS1 is reduced in HCC and higher expression of this transcript is associated with significantly better recurrence-free survival (RFS)^51^. These results suggest that SH3RF3-AS1 may function as an anti-oncogene.

Overall, our results suggest that lncRNAs represent an avenue with great potential for discovering novel therapeutic targets and expanding our understanding of Zika’s molecular signature in tumors. Interestingly, five of the identified lncRNAs have not been explored in the context of Zika-induced oncolysis in neural cancers. Further investigation into the relationship between lncRNAs and Zika’s neuro-oncolytic effect may identify novel mechanisms and targets for adjuvant oncolytic virotherapy in the treatment of GBM.

## ACKNOWLEDGMENTS

We thank the faculty and staff at the Center for Genetic Medicine Research at Children’s National Research Institute for their support.

## AUTHOR CONTRIBUTIONS

YAK, SS, and TM conceived and designed the study and wrote the manuscript. TM, SS, and AH acquired the data and completed the meta-analysis. AH, SS, FT, JN, LH, and YAK designed the validation experiments. AH, FT, and JN contributed and/or performed the validation experiments. SS, AH, LH, TM, and YAK analyzed the results. All authors reviewed and edited the paper. YAK, SS, and TM were responsible for the manuscript and for incorporating feedback and suggested edits.

