## Supplemental Files for "Transcriptomic Meta-analysis Identifies Long Non-Coding RNAs Mediating Zika’s Oncolytic Impact in Glioblastoma Multiforme"

| 1st Author | Manuscript Title | SRA ID | Number of Datasets | Zika Strain(s) | Sample Type | Sequencing Platform |
|---|---|---|---|---|---|---|
| Bonenfant | Asian Zika Virus Isolate Significantly Changes the Transcriptional Profile and Alternative RNA Splicing Events in a Neuroblastoma Cell Line | PRJNA630088 | 2 | PRVABC59<br>MR766 | SH-SY5Y Neuroblastoma cells | Illumina NextSeq 500 |
| Zhu | Zika virus has oncolytic activity against glioblastoma stem cells | PRJNA399336 | 1 | Dakar 41519 | Glioblastoma stem cells (GSCs) and differentiated glioma cells (DGCs) | Illumina HiSeq 3000 |
| Bulstrode | Myeloid cell interferon secretion restricts Zika flavivirus infection of developing and malignant human neural progenitor cells | PRJNA739733 | 1 | PE243 | E22 and E34 Glioblastoma cells | Illumina Novaseq 6000 |

**Table S1. Overview of meta-analysis studies.**

| Gene | Sequence 5' → 3' |
|---|---|
| LINC03032 | Fwd: ACCCATCTTATCCAGGGGCT<br>Rev: AGCATCATGTGGGTGACTGG |
| TIPARP-AS1 | Fwd: AATCAATCACCCCGATCCGC<br>Rev: GTTCTCTCAGGCTCCCGTTC |
| SH3RF3-AS1 | Fwd: CTGCGGGCTACACCTGTC<br>Rev: CTTCCTGCCGCTGGTCTC |
| EMC3-AS1 | Fwd: AGAAAGCCAGATGTCAGGGC<br>Rev: CATGCTGGGGGCTTCACTAA |
| HCG18 | Fwd: CTCCCTCTTTGCCCACTAGC<br>Rev: CACTTGGAGATCCAGGCTCG |
| MELTF-AS1 | Fwd: CAAGTCATTTCAGGGCCCCA<br>Rev: CAGTGCTTCTGAACGCCTCT |

**Table S2. lncRNA primers used for quantitative real-time PCR.**

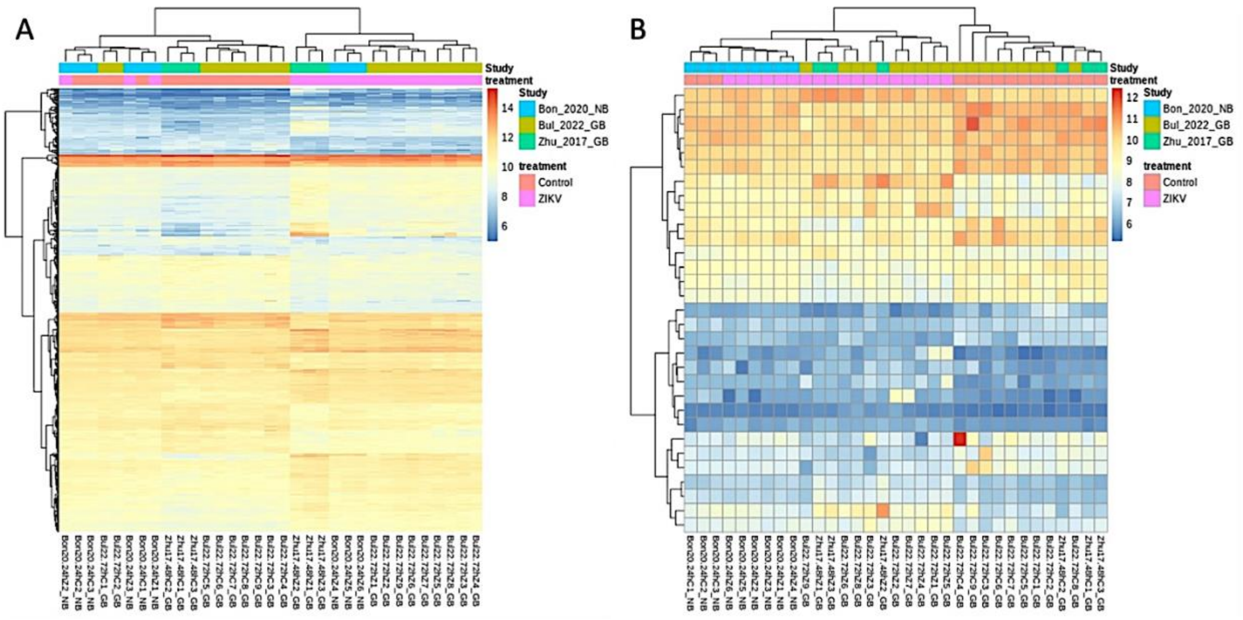


**Figure S1. Heatmaps of coding and non-coding genes.** Heatmaps depicting the transcriptional patterns of differentially expressed coding genes (A) and non-coding genes (B) between ZIKV-infected and control GBM and NBM across the three studies.

A

| Entrez ID | Symbol | Description | Predicted Expression |
|---|---|---|---|
| 7412 | VCAM1 | vascular cell adhesion molecule 1 | Upregulated |
| 3383 | ICAM1 | intercellular adhesion molecule 1 | Upregulated |
| 1649 | DDIT3 | DNA damage inducible transcript 3 | Upregulated |
| 8638 | OASL | 2'-5'-oligoadenylate synthetase like | Upregulated |
| 5971 | RELB | RELB proto-oncogene, NF-kB subunit | Upregulated |
| 2669 | GABRA6 | gamma-aminobutyric acid type A receptor subunit alpha6 | Upregulated |
| 54498 | SMOX | spermine oxidase | Upregulated |
| 90637 | ZFAND2A | zinc finger AN1-type containing 2A | Upregulated |
| 8767 | RIPK2 | receptor interacting serine/threonine kinase 2 | Upregulated |
| 51009 | DERL2 | derlin 2 | Upregulated |
| 54934 | KANSL2 | KAT8 regulatory NSL complex subunit 2 | Upregulated |
| 64782 | AEN | apoptosis enhancing nuclease | Upregulated |
| 9197 | SLC33A1 | solute carrier family 33 member 1 | Upregulated |
| 330 | BIRC3 | baculoviral IAP repeat containing 3 | Upregulated |
| 4189 | DNAJB9 | DnaJ heat shock protein family (Hsp40) member B9 | Upregulated |
| 3726 | JUNB | JunB proto-oncogene, AP-1 transcription factor subunit | Upregulated |
| 64089 | SNX16 | sorting nexin 16 | Upregulated |
| 50484 | RRM2B | ribonucleotide reductase regulatory TP53 inducible subunit M2B | Upregulated |
| 467 | ATF3 | activating transcription factor 3 | Upregulated |
| 27230 | SERP1 | stress associated endoplasmic reticulum protein 1 | Upregulated |
| 3669 | ISG20 | interferon stimulated exonuclease gene 20 | Upregulated |
| 51191 | HERC5 | HECT and RLD domain containing E3 ubiquitin protein ligase 5 | Upregulated |
| 221895 | JAZF1 | JAZF zinc finger 1 | Upregulated |
| 285636 | RIMOC1 | RAB7A interacting MON1-CCZ1 complex subunit 1 | Upregulated |
| 9709 | HERPUD1 | homocysteine inducible ER protein with ubiquitin like domain 1 | Upregulated |
| 4783 | NFIL3 | nuclear factor, interleukin 3 regulated | Upregulated |
| 10140 | TOB1 | transducer of ERBB2, 1 | Upregulated |
| 4791 | NFKB2 | nuclear factor kappa B subunit 2 | Upregulated |
| 26511 | CHIC2 | cysteine rich hydrophobic domain 2 | Upregulated |
| 55254 | TMEM29A | transmembrane protein 39A | Upregulated |
| 4793 | NFKBIB | NFKB inhibitor beta | Upregulated |
| 6236 | RRAD | RRAD, Ras related glycolysis inhibitor and calcium channel regulator | Upregulated |
| 8878 | SQSTM1 | sequestosome 1 | Upregulated |
| 7494 | XBP1 | X-box binding protein 1 | Upregulated |
| 84919 | PPP1R15B | protein phosphatase 1 regulatory subunit 15B | Upregulated |
| 23471 | TRAM1 | translocation associated membrane protein 1 | Upregulated |
| 23764 | MAFF | MAF bZIP transcription factor F | Upregulated |
| 3656 | IRAK2 | interleukin 1 receptor associated kinase 2 | Upregulated |
| 7185 | TRAF1 | TNF receptor associated factor 1 | Upregulated |
| 2908 | NR3C1 | nuclear receptor subfamily 3 group C member 1 | Upregulated |
| 81788 | NUAK2 | NUAK family kinase 2 | Upregulated |
| 7538 | ZFP36 | ZFP36 ring finger protein | Upregulated |
| 3659 | IRF1 | interferon regulatory factor 1 | Upregulated |
| 150274 | HSCB | HscB mitochondrial iron-sulfur cluster cochaperone | Upregulated |
| 9592 | IER2 | immediate early response 2 | Upregulated |
| 4790 | NFKB1 | nuclear factor kappa B subunit 1 | Upregulated |
| 56927 | GPR108 | G protein-coupled receptor 108 | Upregulated |
| 220213 | OTUD1 | OTU deubiquitinase 1 | Upregulated |
| 4792 | NFKBIA | NFKB inhibitor alpha | Upregulated |
| 23092 | ARHGAP26 | Rho GTPase activating protein 26 | Upregulated |
| 27113 | BBC3 | BCL2 binding component 3 | Upregulated |
| 65084 | TMEM135 | transmembrane protein 135 | Upregulated |
| 373 | TRIM23 | tripartite motif containing 23 | Upregulated |
| 6352 | CCL5 | C-C motif chemokine ligand 5 | Upregulated |
| 51307 | FAM53C | family with sequence similarity 53 member C | Upregulated |
| 2181 | ACSL3 | acyl-CoA synthetase long chain family member 3 | Upregulated |
| 51278 | IER5 | immediate early response 5 | Upregulated |
| 51061 | TXNDC11 | thioredoxin domain containing 11 | Upregulated |
| 64651 | CSRNP1 | cysteine and serine rich nuclear protein 1 | Upregulated |
| 83667 | SESN2 | sestrin 2 | Upregulated |
| 3433 | IFIT2 | interferon induced protein with tetratricopeptide repeats 2 | Upregulated |
| 6811 | STX5 | syntaxin 5 | Upregulated |
| 23645 | PPP1R15A | protein phosphatase 1 regulatory subunit 15A | Upregulated |
| 4084 | MXD1 | MAX dimerization protein 1 | Upregulated |
| 9619 | ABCG1 | ATP binding cassette subfamily G member 1 | Upregulated |
| 1263 | PLK3 | polo like kinase 3 | Upregulated |
| 9572 | NR1D1 | nuclear receptor subfamily 1 group D member 1 | Upregulated |
| 694 | BTG1 | BTG anti-proliferation factor 1 | Upregulated |
| 51030 | TVP23B | trans-golgi network vesicle protein 23 homolog B | Upregulated |
| 9095 | TBX19 | T-box transcription factor 19 | Upregulated |
| 10802 | SEC24A | SEC24 homolog A, COPII coat complex component | Upregulated |
| 1326 | MAP3K8 | mitogen-activated protein kinase kinase kinase 8 | Upregulated |
| 8780 | RIOK3 | RIO kinase 3 | Upregulated |
| 55602 | CDKN2AIP | CDKN2A interacting protein | Upregulated |
| 27289 | RND1 | Rho family GTPase 1 | Upregulated |
| 7358 | UGDH | UDP-glucose 6-dehydrogenase | Upregulated |
| 9546 | APBA3 | amyloid beta precursor protein binding family A member 3 | Upregulated |
| 96459 | FNIP1 | folliculin interacting protein 1 | Upregulated |
| 64764 | CREB3L2 | cAMP responsive element binding protein 3 like 2 | Upregulated |
| 4616 | GADD45B | growth arrest and DNA damage inducible beta | Upregulated |
| 3717 | JAK2 | Janus kinase 2 | Upregulated |
| 3638 | INSIG1 | insulin induced gene 1 | Upregulated |
| 1294 | COL7A1 | collagen type VII alpha 1 chain | Upregulated |
| 6347 | CCL2 | C-C motif chemokine ligand 2 | Upregulated |
| 6319 | SCD | stearoyl-CoA desaturase | Upregulated |
| 1052 | CEBPD | CCAAT enhancer binding protein delta | Upregulated |
| 2643 | GCH1 | GTP cyclohydrolase 1 | Upregulated |
| 43 | ACHE | acetylcholinesterase (Yt blood group) | Upregulated |
| 51726 | DNAJB11 | DnaJ heat shock protein family (Hsp40) member B11 | Upregulated |
| 3309 | HSPA5 | heat shock protein family A (Hsp70) member 5 | Upregulated |
| 7873 | MANF | mesencephalic astrocyte derived neurotrophic factor | Upregulated |
| 5228 | PGF | placental growth factor | Upregulated |
| 8553 | BHLHE40 | basic helix-loop-helix family member e40 | Upregulated |
| 9601 | PDIA4 | protein disulfide isomerase family A member 4 | Upregulated |
| 10525 | HYOU1 | hypoxia up-regulated 1 | Upregulated |
| 79174 | CRELD2 | cysteine rich with EGF like domains 2 | Upregulated |
| 23753 | SDF2L1 | stromal cell derived factor 2 like 1 | Upregulated |
| 9719 | ADAMTSL2 | ADAMTS like 2 | Upregulated |
| 55715 | DOK4 | docking protein 4 | Downregulated |
| 10220 | GDF11 | growth differentiation factor 11 | Downregulated |
| 4001 | LMNB1 | lamin B1 | Downregulated |
| 400506 | KNOP1 | lysine rich nucleolar protein 1 | Downregulated |
| 85449 | KIAA1755 | KIAA1755 | Downregulated |
| 51280 | GOLM1 | golgi membrane protein 1 | Downregulated |
| 2583 | B4GALNT1 | beta-1,4-N-acetyl-galactosaminyltransferase 1 | Downregulated |
| 56135 | PCDHAC1 | protocadherin alpha subfamily C, 1 | Downregulated |
| 1463 | NCAN | neurocan | Downregulated |
| 2137 | EXTL3 | exostosin like glycosyltransferase 3 | Downregulated |
| 56098 | PCDHGC4 | protocadherin gamma subfamily C, 4 | Downregulated |
| 23240 | TMEM131L | transmembrane 131 like | Downregulated |
| 83879 | CDCA7 | cell division cycle associated 7 | Downregulated |
| 5764 | PTN | pleiotrophin | Downregulated |
| 152002 | XXYLT1 | xyloside xylosyltransferase 1 | Downregulated |
| 6941 | TCF19 | transcription factor 19 | Downregulated |
| 7112 | TMPO | thymopoietin | Downregulated |
| 899 | CCNF | cyclin F | Downregulated |
| 57462 | MYORG | myogenesis regulating glycosidase | Downregulated |
| 79888 | LPCAT1 | lysophosphatidylcholine acyltransferase 1 | Downregulated |
| 9498 | SLC4A8 | solute carrier family 4 member 8 | Downregulated |
| 150946 | GAREM2 | GRB2 associated regulator of MAPK1 subtype 2 | Downregulated |
| 2969 | GTF2I | general transcription factor Iii | Downregulated |
| 25758 | KIAA1549L | KIAA1549 like | Downregulated |
| 9917 | FAM20B | FAM20B glycosaminoglycan xylosylkinase | Downregulated |
| 51435 | SCARA3 | scavenger receptor class A member 3 | Downregulated |
| 65998 | ZFTA | zinc finger translocation associated | Downregulated |
| 57556 | SEMA6A | semaphorin 6A | Downregulated |
| 79966 | SCD5 | stearoyl-CoA desaturase 5 | Downregulated |
| 50512 | PODXL2 | podocalyxin like 2 | Downregulated |
| 10236 | HNRNPR | heterogeneous nuclear ribonucleoprotein R | Downregulated |
| 2899 | GRIK3 | glutamate ionotropic receptor kainate type subunit 3 | Downregulated |
| 92370 | PXYLP1 | 2-phosphoxylose phosphatase 1 | Downregulated |
| 54058 | C21orf58 | chromosome 21 open reading frame 58 | Downregulated |
| 9315 | NREP | neuronal regeneration related protein | Downregulated |
| 8914 | TIMELESS | timeless circadian regulator | Downregulated |
| 5098 | PCDHGC3 | protocadherin gamma subfamily C, 3 | Downregulated |
| 5738 | PTGFRN | prostaglandin F2 receptor inhibitor | Downregulated |
| 3488 | IGFBP5 | insulin like growth factor binding protein 5 | Downregulated |
| 84913 | ATOH8 | atonal bHLH transcription factor 8 | Downregulated |
| 9806 | SPOCK2 | SPARC (osteonectin), cwcv and kazal like domains proteoglycan 2 | Downregulated |
| 7168 | TPM1 | tropomyosin 1 | Downregulated |
| 1844 | DUSP2 | dual specificity phosphatase 2 | Downregulated |
| 338382 | RAB7B | RAB7B, member RAS oncogene family | Downregulated |
| 27253 | PCDH17 | protocadherin 17 | Downregulated |
| 115207 | KCTD12 | potassium channel tetramerization domain containing 12 | Downregulated |

B

| Entrez ID | Gene Type | Symbol | Description | Predicted Expression in GBM |
|---|---|---|---|---|
| 124909347 | ncRNA | LOC124909347 | Uncharacterized | Upregulated |
| 124907970 | ncRNA | LOC124907970 | Uncharacterized | Upregulated |
| 105372436 | ncRNA | LOC105372436 | Uncharacterized | Upregulated |
| 441094 | ncRNA | NR2F1-AS1 | NR2F1 antisense RNA 1 | Upregulated |
| 105370361 | ncRNA | LINC03032 | Long intergenic non-protein coding RNA 3032 | Upregulated |
| 100287227 | ncRNA | TIPARP-AS1 | TIPARP antisense RNA 1 | Upregulated |
| 100287216 | ncRNA | SH3RF3-AS1 | SH3RF3 antisense RNA 1 | Upregulated |
| 107983990 | ncRNA | LOC107983990 | Uncharacterized | Upregulated |
| 105370449 | ncRNA | LOC105370449 | Uncharacterized | Upregulated |
| 100996712 | miscRNA | SRGAP2D | Pseudogene | Upregulated |
| 644961 | miscRNA | ACTG1P20 | Pseudogene | Upregulated |
| 100132057 | miscRNA | PDE4DIPP6 | Pseudogene | Upregulated |
| 124904006 | ncRNA | LOC124904006 | Uncharacterized | Upregulated |
| 105375914 | ncRNA | LOC105375914 | Uncharacterized | Upregulated |
| 100132147 | ncRNA | LINC01783 | long intergenic non-protein coding RNA 1783 | Upregulated |
| 284600 | ncRNA | LOC284600 | Uncharacterized | Downregulated |
| 155400 | miscRNA | NSUN5P1 | Pseudogene | Downregulated |
| 105375304 | ncRNA | SNX10-AS1 | SNX10 antisense RNA 1 | Downregulated |
| 100129482 | miscRNA | ZNF37BP | Pseudogene | Downregulated |
| 130872 | miscRNA | AHSA2P | Pseudogene | Downregulated |
| 100132249 | ncRNA | LOC100132249 | Uncharacterized | Downregulated |
| 55073 | miscRNA | LRRC37A4P | Pseudogene | Downregulated |
| 100288152 | ncRNA | SLC9A3-AS1 | SLC9A3 antisense RNA 1 | Downregulated |
| 442075 | ncRNA | EMC3-AS1 | EMC3 antisense RNA 1 | Downregulated |
| 414777 | ncRNA | HCG18 | HLA complex group 18 | Downregulated |
| 100507057 | ncRNA | MELTF-AS1 | MELTF antisense RNA 1 | Downregulated |
| 100506421 | ncRNA | PANTR1 | POU3F3 adjacent non-coding transcript 1 | Downregulated |
| 29774 | miscRNA | POM121L9P | Pseudogene | Downregulated |
| 503638 | miscRNA | DUXAP9 | Pseudogene | Downregulated |
| 401303 | miscRNA | ZNF815P | Pseudogene | Downregulated |
| 100532732 | ncRNA | MSH5-SAPCD1 | MSH5-SAPCD1 readthrough (NMD candidate) | Downregulated |

**Fig S2. Differentially Expressed Genes.** Lists of the coding genes (A) significantly dysregulated in at least three out of the four datasets and non-coding genes (B) consistently dysregulated between the two GBM studies.
